# Backward compatibility of whole genome sequencing data with MLVA typing using a new *MLVAtype* shiny application: the example of *Vibrio cholerae*

**DOI:** 10.1101/663138

**Authors:** Jérôme Ambroise, Léonid M. Irenge, Jean-François Durant, Bertrand Bearzatto, Godfrey Bwire, O. Colin Stine, Jean-Luc Gala

## Abstract

Multiple-Locus Variable Number of Tandem Repeats (VNTR) Analysis (MLVA) is widely used by laboratory-based surveillance networks for subtyping pathogens causing foodborne and water-borne disease outbreaks. However, Whole Genome Sequencing (WGS) has recently emerged as the new more powerful reference for pathogen subtyping, making a data conversion method necessary which enables the users to compare the MLVA identified by either method. The *MLVAType* shiny application was designed to extract MLVA profiles from WGS data while ensuring backward compatibility with traditional MLVA typing methods.

To test and validate the *MLVAType* algorithm, WGS-derived MLVA profiles of nineteen *Vibrio cholerae* isolates from Democratic Republic of the Congo (n=9) and Uganda (n=10) were compared to MLVA profiles generated by microchip electrophoresis (Bioanalyzer Agilent 2100), GeneScan analysis, and Sanger sequencing as the reference method. Unlike amplicon-size derived MLVA profiles, results obtained by Sanger sequencing and WGS were totally concordant. However, the latter were affected by censored estimations whose percentage was inversely proportional to the k-mer parameter used during genome assembly. With a k-mer of 127, less than 15% estimation of *V. cholerae* VNTR was censored. Preventing censored estimation was only achievable when using a longer k-mer size (*i*.*e*. 175), which is not proposed in the SPAdes v.3.13.0 software.

*In silico* analysis showed that this limitation does not apply to other microbial species (*e*.*g. Mycobacterium, Streptococcus, Staphylococcus*, and *Pseudomonas*) characterized by smaller lengths of motif repeats. As NGS read lengths and qualities tend to increase with time, one may expect the increase of k-mer size in a near future. Using *MLVAType* application with a longer k-mer size will then efficiently retrieve MLVA profiles from WGS data while avoiding censored estimation irrespective of the microbial species.

**Author summary:** Next Generation Sequencing (NGS) has emerged as a powerful high throughput genomic approach enabling the Whole Genome Sequence (WGS) of pathogens to be assembled in a relatively short time. A major advantage of WGS, compared to traditional genotypic identification and typing methods, is its ability to generate data that can be exploited *in silico* for multiple bacterial tests including accurate subtyping, determination of genetic relatedness, and characterization of virulence and antimicrobial resistance determinants. Accordingly, WGS is now rapidly replacing traditional methods like Multi-Locus Variable Number of Tandem Repeats Analysis (MLVA) that has long been used in the public health sector for laboratory-based surveillance of pathogens and outbreak response. While these missions require maintenance of data comparability within networks, the lack of backward compatibility between WGS-derived and traditional MLVA methods is a well-recognized issue. As illustrated here with *Vibrio cholerae* isolates from DRC and Uganda, the *MLVAType* software application analyzes WGS data to generate MLVA profiles that are identical to those determined with traditional typing. Interestingly, this tool has also the potential to extract MLVA profiles from any bacterial genome that are characterized by a small number of tandem repeats, *e*.*g. Streptococcus, Staphylococcus, Pseudomonas*, and *Mycobacterium* species. This restriction can be lifted if subsequences of length k, called k-mers, are longer than what is currently proposed by genome assembly algorithm like SPAdes v.3.13.0.

## Introduction

Rapid molecular typing of pathogens associated with human and animal diseases has proven instrumental in the surveillance and control of infectious diseases [1, 2]. Pulsed field gel-electrophoresis (PFGE), which was long considered as the gold standard for molecular typing of pathogens associated with outbreaks, has been superseded by Multi-Locus Sequence Typing (MLST) or Multi-Locus Variable Number of Tandem Repeats (VNTR) Analysis (MLVA), and more recently by Whole Genome Sequencing (WGS). Mounting evidence supports the use of the latter method as a new gold standard for pathogen subtyping [3].

However, unlike MLVA, WGS analysis requires a specific expertise in bioinformatics and is not yet affordable in all developing countries where highly pathogenic diseases would make it the most useful and such a method would be the most needed. One can therefore be sure that MLVA and WGS subtyping will coexist in years to come, making necessary a methodology enabling end-users (*i*.*e*. researchers, clinicians, microbiologist and epidemiologists) to compare respective results.

Accordingly, *in silico* methods which extract low-throughput typing results (e.g., MLST or MLVA) from WGS data should be developed to enable users to compare subtyping results irrespective of the methodology and time of data acquisition. Both parameters are important when WGS data need to be compared with data generated before the WGS era.

Whereas the number of tandem repeats at different VNTR loci may theoretically be retrieved from WGS data, like currently done when extracting MLST from WGS, this was practically not considered feasible with MLVA because of a lack of accuracy of genomes assembly derived from Next Generation Sequencing (NGS) short reads [4]. Limited backward compatibility of WGS with MLVA is indeed notoriously due to failure to correctly assemble repetitive regions assessed by MLVA [5]. When analyzing NGS data, the first step generally consists in assembling reads into longer contiguous sequences (contigs), which can then be interrogated using BLAST or other search algorithms. The production of high quality assemblies using bacterial genome assembler such as SPAdes [6] requires quality filtering and optimization of different parameters including k-mer size.

In the current paper, we describe a new tool (named *MLVAtype*) which enables users to extract MLVA profiles from WGS data. We tuned the k-mer parameter used during genome assembly in order to assess its impact on the performance and limitations of *MLVAtype*. As a proof of concept, this new tool was applied on draft genomes of *V. cholerae* isolates associated with cholera outbreaks in two bordering countries, *i*.*e*. the Democratic Republic of the Congo (DRC) and Uganda. Additionally, the feasibility of using *MLVAtype* to type other bacterial pathogens was deduced from MLVA profiles (number of tandem repeats) downloaded from a public database.

## Materials and methods

### Sample description

Nine *V. cholerae* isolates were selected from a collection of isolates characterized in a recent study conducted between 2014 and 2017 in the DRC [7]. In addition, ten *V. cholerae* isolates collected between 2014 and 2016 in Uganda by G. Bwire and colleagues were selected based on their published data [8].

### Amplicon size-versus Sanger-derived MLVA typing

Amplicon size determination of DRC isolates was carried out using an Agilent 2100 Bioanalyzer in conjunction with the DNA 1000 and the High Sensitivity DNA Labchips Kits (Agilent Technologies, Waldbronn, Germany) according to the manufacturer’s recommendations. In parallel, amplicons were sequenced on both strands on the ABI 3130 GA, using the BigDye Terminator v1.1 cycle sequencing kit (Applied Biosystems, USA). Motif repeats were counted manually and translated into MLVA profiles.

For Ugandan isolates, amplicon size determination was retrieved from published data [8] where fluorescently labeled amplified products were separated using a 3730xl Automatic Sequencer with the size determined from internal lane standards (LIZ600) by the GeneScan program (Applied Biosystems, Foster City, CA).

When MLVA profiles were generated according to the method proposed by Kendall et al. [9], the formula had to be modified to better fit the sequence length of the motif and the position of the primers (Table 1). It is of note that, the original calculation formula was used for the VC0283 motif of Ugandan isolates but with modified primers positions (data not shown).

**Table 1.**
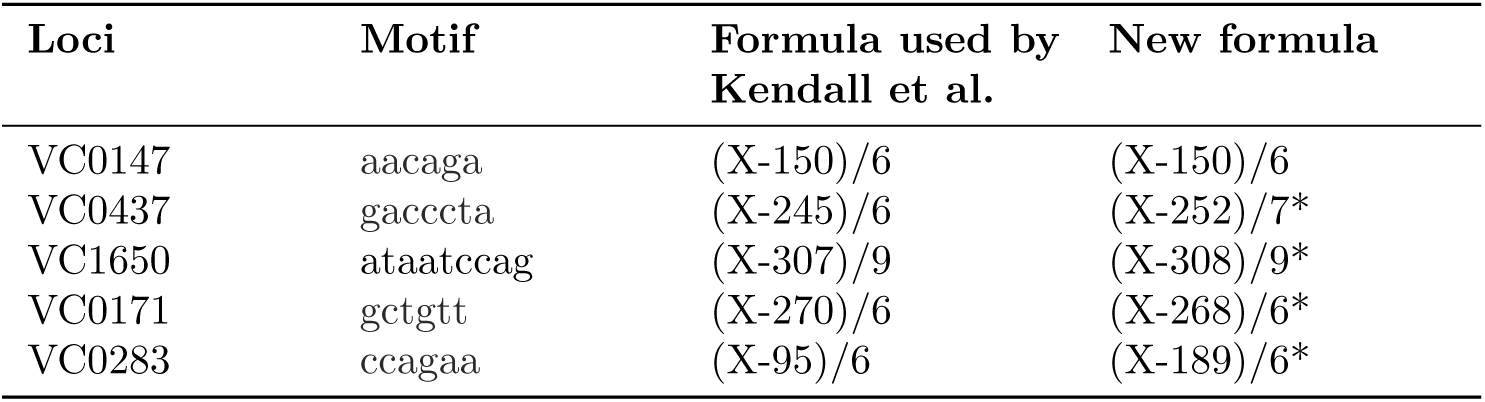
Formula used in the current study to compute the number of tandem repeats from the amplicon size. *: Modifications introduced in the new formula

## WGS-derived MLVA typing

### WGS and short reads assembly

Whole genome assemblies of selected *V. cholerae* isolates, 9 from DRC and 10 from Uganda, were generated from paired-end 300 and 150 nt long reads, respectively. In brief, genomic DNA from DRC isolates was simultaneously fragmented and tagged with sequencing adapters in a single step using Nextera transposome (Nextera XT DNA Library Preparation Kit, Illumina, San Diego, CA, USA). Tagged DNA was then amplified with a 12-cycle polymerase chain reaction (PCR), cleaned up with AMPure beads, and subsequently loaded on a MiSeq paired-end 2 × 300 nt (MiSeq reagent kit V3 (600 cycles) sequence run. Raw genomic data were submitted to the European Nucleotide Archive (ENA, http://www.ebi.ac.uk/ena), and are available under study accession number ERP114722. For Ugandan *V. cholerae* isolates, data which were retrieved from the previous publication [7], were obtained as follows: libraries for Illumina sequencing were prepared from DNA fragmented with Covaris E210 (Covaris, Wolburn, MA) using the KAPA High Throughput Library Preparation Kit (Millipore-Sigma, St. Louis MO). The libraries were enriched and barcoded in ten cycles of PCR amplification with primers containing an index sequence. Subsequently, the libraries were sequenced using a 150 nt paired-end run on an Illumina HiSeq2500 (Illumina, San Diego, CA). Raw genomic data were submitted to Sequence Read Archive (SRA, https://www.ncbi.nlm.nih.gov/sra) under study accession number PRJNA439310.

WGS data were assembled into contigs using SPAdes v.3.13.0 [6] and a range of k-mer sizes of 55, 77, 99, 127, and 175-mers. Testing a k-mer value of 175, which has a longer size than what is proposed in the SPAdes v.3.13.0 software, required editing the software source code. It has to be kept in mind that using this longer k-mer size is only possible with reads of sufficient length. Accordingly, a k-mer size of 175 was only used with 300 nt reads from DRC isolates, but not with 150 nt reads from Ugandan isolates. For each sample, k-mer size and locus, the number of tandem repeats was extracted from the assembled draft genome using the *MLVAtype* algorithm.

### *MLVAtype* algorithm

The *MLVAtype* algorithm processes each motif separately. It requires several inputs including a draft genome, the size of the k-mer (*i*.*e*. k-mer parameter) used during genome assembly, and the nucleotide sequence of the motif. The algorithm returns the number of tandem repeats using the following steps: first, small contigs (< 2000 nt) are removed from the draft genome. Second, the vcountPattern function from the Biostring R package is used to count the number of occurrences that a single (j=1) motif is detected within the draft genome. Then the same computation is iteratively performed with an increasing number (j=2, 3 .., k) of tandem repeats. This iterative process is performed until there is only one occurrence of the k tandem repeats. Finally, the maximum value of j (*i*.*e*. k) is compared to the maximum number of tandem repeats (MNTR) that can be included in a specified k-mer (Fig 1 and Table 2). If k ≥ MNTR, the estimation of the number of tandem repeats is set to MNTR and considered as right-censored (*i*.*e*. ≥MNTR). If k < MNTR, the estimation of the number of tandem repeats is set to k. The same process is applied on each motif in order to extract the complete MLVA profile from the genome assembly.

**Table 2.** Maximum number of tandem repeats in a k-mer

| Loci | motif length | k-mer = 33 | k-mer = 55 | k-mer = 77 | k-mer = 99 | k-mer = 127 | k-mer = 175 |
| --- | --- | --- | --- | --- | --- | --- | --- |
| VC0147 | 6 | 5 | 9 | 12 | 16 | 21 | 29 |
| VC0437 | 7 | 4 | 7 | 11 | 14 | 18 | 25 |
| VC1650 | 9 | 3 | 6 | 8 | 11 | 14 | 19 |
| VCA0171 | 6 | 5 | 9 | 12 | 16 | 21 | 29 |
| VCA0283 | 6 | 5 | 9 | 12 | 16 | 21 | 29 |

**Fig 1.**
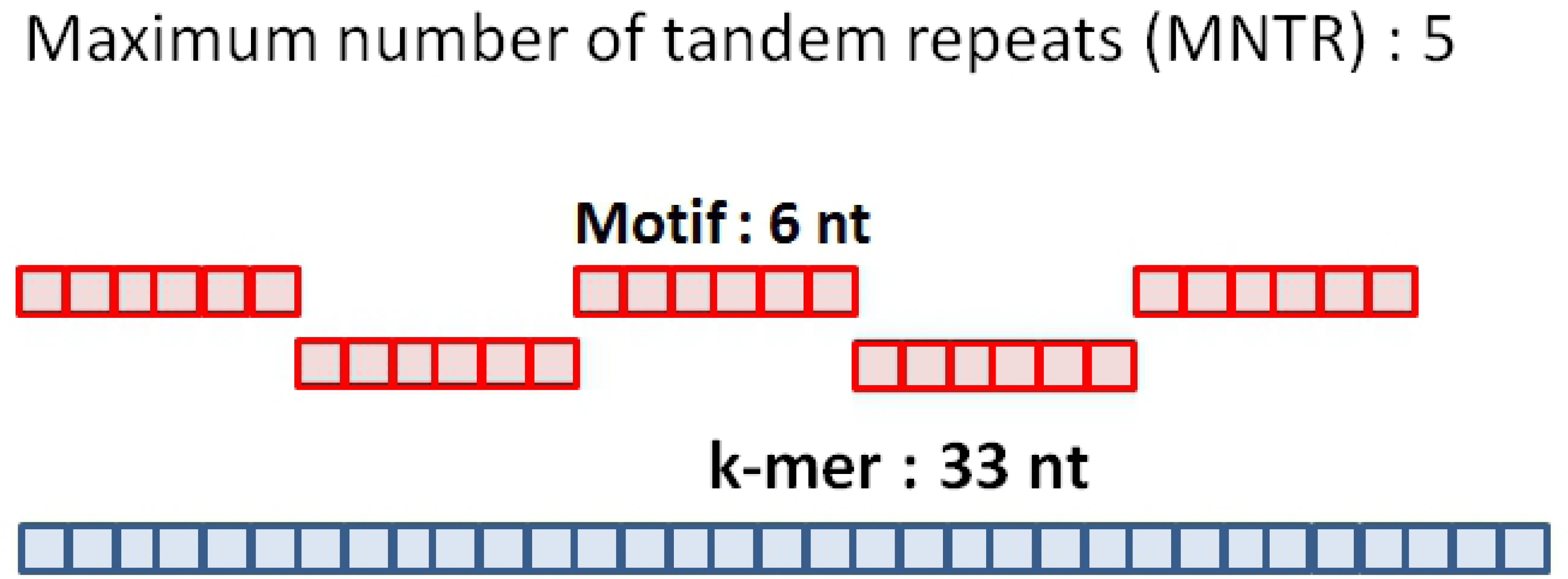
Maximum number of tandem repeats (MNTR). Example of the maximum number of a 6 nt motif included in a k-mer of 33 nt.

### *MLVAtype* shiny application

The *MLVAtype* algorithm was implemented in an R shiny application which is freely available at https://ucl-irec-ctma.shinyapps.io/NGS-MLVA-TYPING/. This application enables the user to upload a list of draft genomes, the nucleotide sequences of the motifs, and the value of the k-mer which was used to build the assembly, including a k-mer size selectable after modification of the SPAdes v.3.13.0 source code (Fig 2). The application provides a table with the number of tandem repeats that was found for each motif in the corresponding genomes.

**Fig 2.**
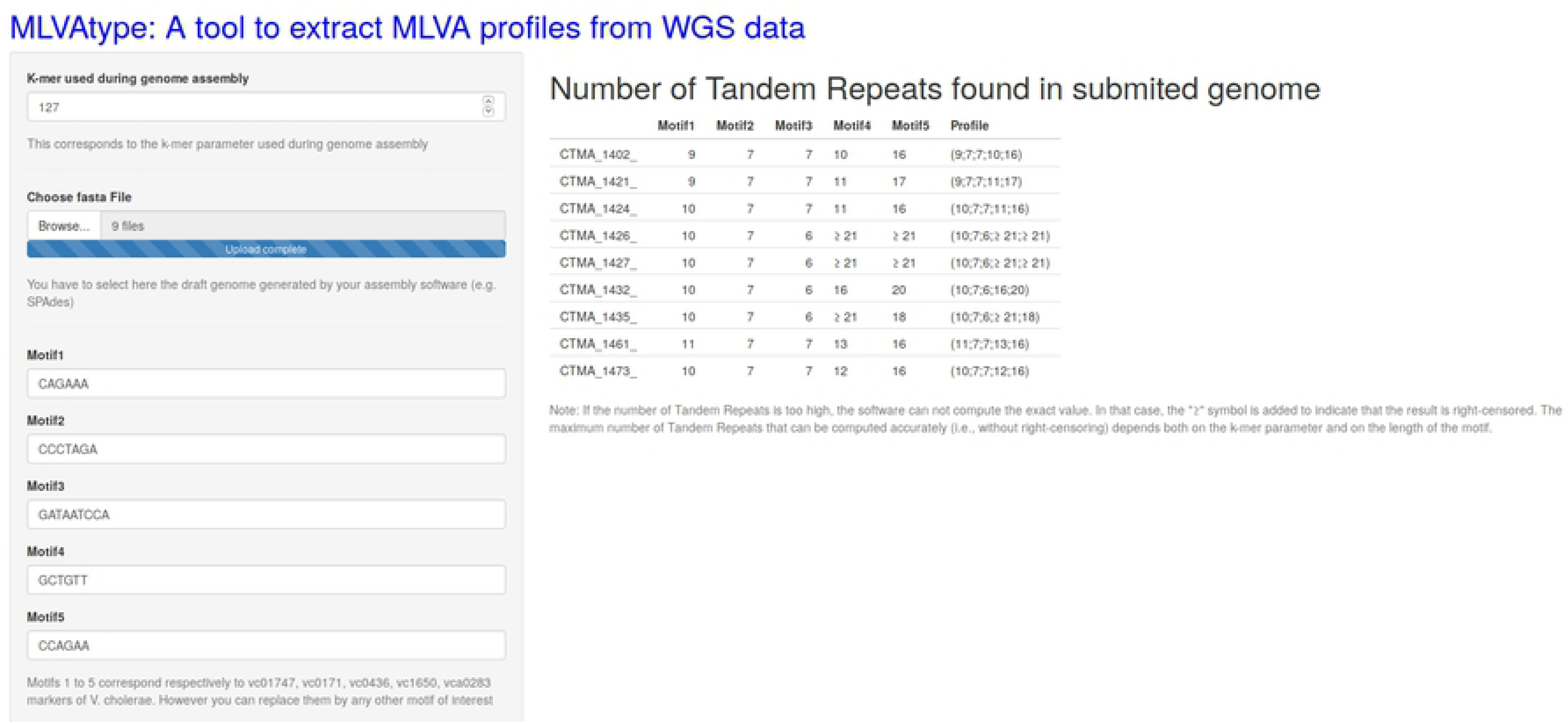
Screenshot of the *MLVAtype* shiny application. Nine *V. cholerae* genomes from DRC were uploaded and processed with this application.

## Results

### Amplicon-size-derived and Sanger-derived MLVA typing

Amplicon size- and Sanger-derived MLVA profiles from DRC and Uganda are reported in Table 3. Considering the high quality of the Sanger sequences, the number of tandem repeats extracted from these sequences were considered as a gold-standard in the current study. As expected, amplicon-size-derived results did not perfectly correlate with those obtained with Sanger sequencing. Furthermore, the proportion of mismatches was much higher when amplicon size was determined with the Bioanalyser than with the GeneScan program (*i*.*e*., 32/45 for DRC isolates versus 8/50 for Ugandan isolates).

**Table 3.** MLVA profiles consisting in the number of tandem repetition for each loci (VC0147, VC0437, VC1650, VCA0171, and VCA0283) extracted from amplicon sizes and from Sanger sequences. *Amplicon-size was obtained using the Bioanalyzer and GeneScan program for DRC and Ugandan isolates, respectively (cf methods). Mismatches are indicated in bold.

| Country | Isolate | Sanger-derived MLVA profile | Amplicon-size derived MLVA profile* |
| --- | --- | --- | --- |
| DRC | CTMA-1402 | (9,7,7,10,16) | (**8**,5,7,**11**,14) |
| DRC | CTMA-1421 | (9,7,7,11,17) | (**8**,5,7,11,**14**) |
| DRC | CTMA-1424 | (10,7,7,11,16) | (**9**,5,7,11,**13**) |
| DRC | CTMA-1426 | (10,7,6,24,21) | (**9**,5,7,**23**,18) |
| DRC | CTMA-1427 | (10,7,6,24,21) | (**9**,5,6,**23**,18) |
| DRC | CTMA-1432 | (10,7,6,16,20) | (**9**,5,6,16,**17**) |
| DRC | CTMA-1435 | (10,7,6,23,18) | (**9**,5,6,**22**,15) |
| DRC | CTMA-1461 | (11,7,7,13,16) | (**9**,5,7,13,**13**) |
| DRC | CTMA-1473 | (10,7,7,12,16) | (**8**,5,7,12,**13**) |
| Uganda | UG010 | (9;3;7;21;26) | (9,**2**,7,21,26) |
| Uganda | UG020 | (9;3;7;21;27) | (9,**2**,7,21,27) |
| Uganda | UG026 | (9;3;7;21;28) | (9,**2**,7,21,28) |
| Uganda | UG040 | (10;7;7;9;17) | (10,7,7,9,17) |
| Uganda | UG042 | (8;7;7;10;21) | (8,**6**,7,10,21) |
| Uganda | UG046 | (8;7;7;11;21) | (8,**6**,7,11,21) |
| Uganda | UG054 | (8;7;7;10;21) | (**5**,7,7,10,21) |
| Uganda | UG060 | (10;7;7;9;17) | (10,**6**,7,9,17) |
| Uganda | UG071 | (10;7;7;8;18) | (10,7,7,8,18) |
| Uganda | UG086 | (10;7;7;9;18) | (10,**6**,7,9,18) |

### WGS-derived MLVA typing

WGS-derived MLVA profiles obtained using the *MLVAtype* algorithm from draft genomes assembled using SPAdes and each value of the k-mer parameter are reported for DRC and Ugandan isolates (Table 4). All estimates of number of tandem repeats appeared to be perfectly concordant with Sanger-derived typing values. However, the k-mer parameter directly impacts the number of right-censored (*i*.*e*., ≥) estimations. When a k-mer of 127 was used to assemble reads from DRC and Ugandan isolates, only 5 and 9 right-censored estimations of tandem repeat numbers were produced, respectively. Altogether, WGS-based method with a k-mer of 127 generated a percentage of correct estimations similar to MLVA typing using the GeneScan program, and caused no incorrect estimation (Fig 3). Interestingly, WGS-derived MLVA profiles extracted from 175-mers DRC assemblies were perfectly concordant with Sanger-derived typing results with no censored estimation.

**Table 4.** WGS-derived MLVA profiles extracted from DRC and Ugandan genomes assembled with various values of the k-mer parameter and compared to the gold-standard (*i*.*e*. Sanger-derived MLVA profile). *: Maximum Number of Tandem Repeats (MNTR) in the corresponding k-mer. n/a: not applicable. Read lengths obtained with DRC and Ugandan isolates were 300 and 150 nt, respectively.

| Country | Isolate | WGS-derived<br>k-mer = 55<br>(9,7,6,9,9)* | WGS-derived<br>k-mer = 77<br>(12,11,8,12,12)* | WGS-derived<br>k-mer = 99<br>(16,14,11,16,16)* | WGS-derived<br>k-mer = 127<br>(21,18,14,21,21)* | WGS-derived<br>k-mer = 175<br>(29,25,19,29,29)* | Sanger-derived MLVA<br>profile |
| --- | --- | --- | --- | --- | --- | --- | --- |
| DRC | CTMA-1402 | (≥9;≥7;≥6;≥9;≥9) | (9;7;7;10;≥12) | (9;7;7;10;≥16) | (9;7;7;10;16) | (9;7;7;10;16) | (9,7,7,10,16) |
| DRC | CTMA-1421 | (≥9;≥7;≥6;≥9;≥9) | (9;7;7;11;≥12) | (9;7;7;11;≥16) | (9;7;7;11;17) | (9;7;7;11;17) | (9,7,7,11,17) |
| DRC | CTMA-1424 | (≥9;≥7;≥6;≥9;≥9) | (10;7;7;11;≥12) | (10;7;7;11;≥16) | (10;7;7;11;16) | (10;7;7;11;16) | (10,7,7,11,16) |
| DRC | CTMA-1426 | (≥9;≥7;≥6;≥9;≥9) | (10;7;6;≥12;≥12) | (10;7;6;≥16;≥16) | (10;7;6;≥21;≥21) | (10;7;6;24;21) | (10,7,6,24,21) |
| DRC | CTMA-1427 | (≥9;≥7;≥6;≥9;≥9) | (10;7;6;≥12;≥12) | (10;7;6;≥16;≥16) | (10;7;6;≥21;≥21) | (10;7;6;24;21) | (10,7,6,24,21) |
| DRC | CTMA-1432 | (≥9;≥7;≥6;≥9;≥9) | (10;7;6;≥12;≥12) | (10;7;6;≥16;≥16) | (10;7;6;16;20) | (10;7;6;16;20) | (10,7,6,16,20) |
| DRC | CTMA-1435 | (≥9;≥7;≥6;≥9;≥9) | (10;7;6;≥12;≥12) | (10;7;6;≥16;≥16) | (10;7;6;≥21;18) | (10;7;6;23;18) | (10,7,6,23,18) |
| DRC | CTMA-1461 | (≥9;≥7;≥6;≥9;≥9) | (11;7;7;≥12;≥12) | (11;7;7;13;≥16) | (11;7;7;13;16) | (11;7;7;13;16) | (11,7,7,13,16) |
| DRC | CTMA-1473 | (≥9;≥7;≥6;≥9;≥9) | (10;7;7;≥12;≥12) | (10;7;7;12;≥16) | (10;7;7;12;16) | (10;7;7;12;16) | (10,7,7,12,16) |
| Uganda | UG010 | (≥9;3;≥6;≥9;≥9) | (9;3;7;≥12;≥12) | (9;3;7;≥16;≥16) | (9;3;7;≥21;≥21) | n/a | (9;3;7;21;26) |
| Uganda | UG020 | (≥9;3;≥6;≥9;≥9) | (9;3;7;≥12;≥12) | (9;3;7;≥16;≥16) | (9;3;7;≥21;≥21) | n/a | (9;3;7;21;27) |
| Uganda | UG026 | (≥9;3;≥6;≥9;≥9) | (9;3;7;≥12;≥12) | (9;3;7;≥16;≥16) | (9;3;7;≥21;≥21) | n/a | (9;3;7;21;28) |
| Uganda | UG040 | (≥9;≥7;≥6;≥9;≥9) | (10;7;7;9;≥12) | (10;7;7;9;≥16) | (10;7;7;9;17) | n/a | (10;7;7;9;17) |
| Uganda | UG042 | (8;≥7;≥6;≥9;≥9) | (8;7;7;10;≥12) | (8;7;7;10;≥16) | (8;7;7;10;≥21) | n/a | (8;7;7;10;21) |
| Uganda | UG046 | (8;≥7;≥6;≥9;≥9) | (8;7;7;11;≥12) | (8;7;7;11;≥16) | (8;7;7;11;≥21) | n/a | (8;7;7;11;21) |
| Uganda | UG054 | (8;≥7;≥6;≥9;≥9) | (8;7;7;10;≥12) | (8;7;7;10;≥16) | (8;7;7;10;≥21) | n/a | (8;7;7;10;21) |
| Uganda | UG060 | (≥9;≥7;≥6;≥9;≥9) | (10;7;7;9;≥12) | (10;7;7;9;≥16) | (10;7;7;9;17) | n/a | (10;7;7;9;17) |
| Uganda | UG071 | (≥9;≥7;≥6;8;≥9) | (10;7;7;8;≥12) | (10;7;7;8;≥16) | (10;7;7;8;18) | n/a | (10;7;7;8;18) |
| Uganda | UG086 | (≥9;≥7;≥6;≥9;≥9) | (10;7;7;9;≥12) | (10;7;7;9;≥16) | (10;7;7;9;18) | n/a | (10;7;7;9;18) |

**Fig 3.**
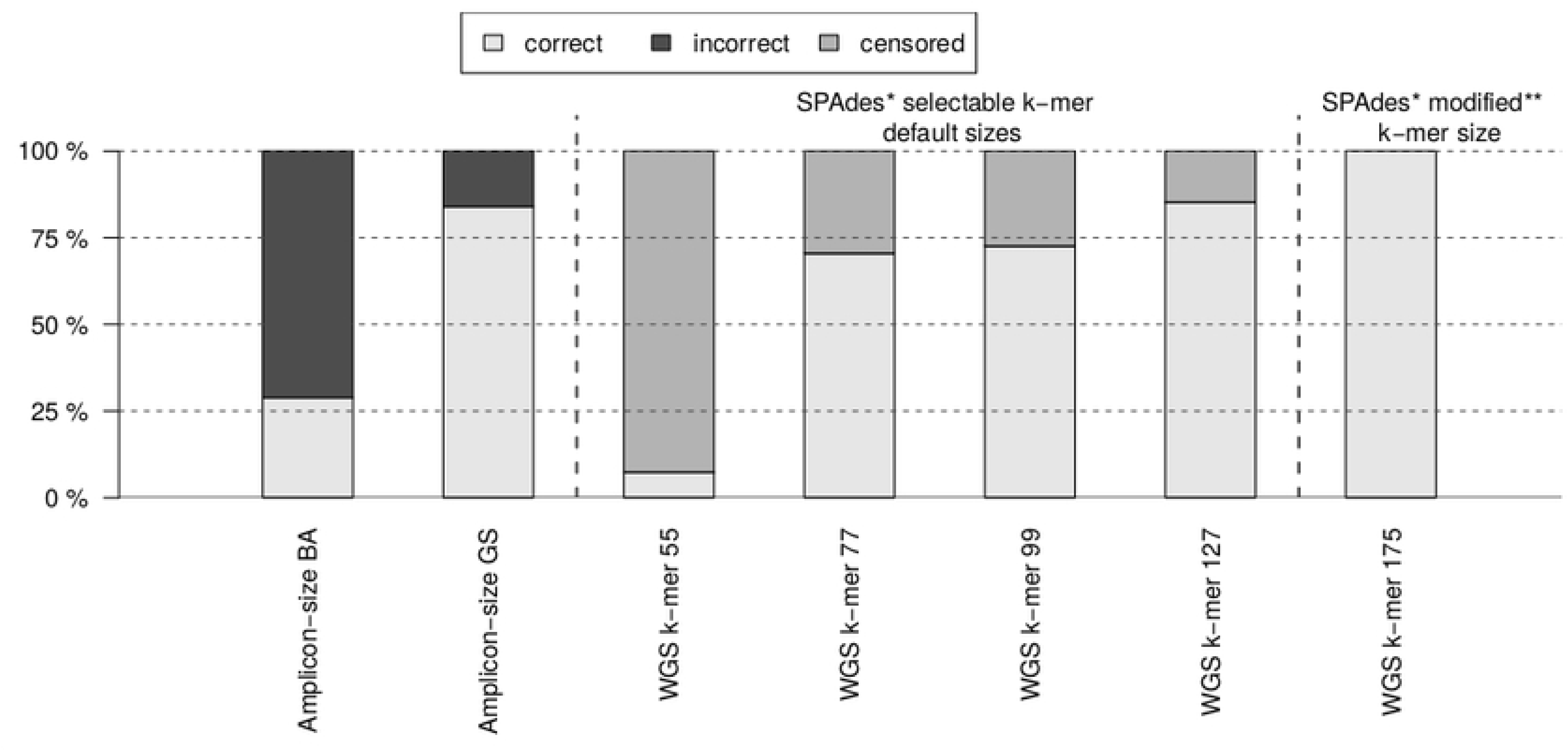
Percentage of correct, incorrect, and censored estimation of the number of tandem repeats. Each percentage is produced by either amplicon size (BA: Bioanalyzer, GS: GeneScan) or WGS-based approaches; Sanger sequencing results were used as the reference values. *: SPAdes v.3.13.0. **: After modification of SPAdes v.3.13.0 source code.

It is worth noting that a longer k-mer size also improved the quality of genome assemblies as illustrated by QUAST [10] with lower numbers of contigs and larger N50 values (Fig 4).

**Fig 4.**
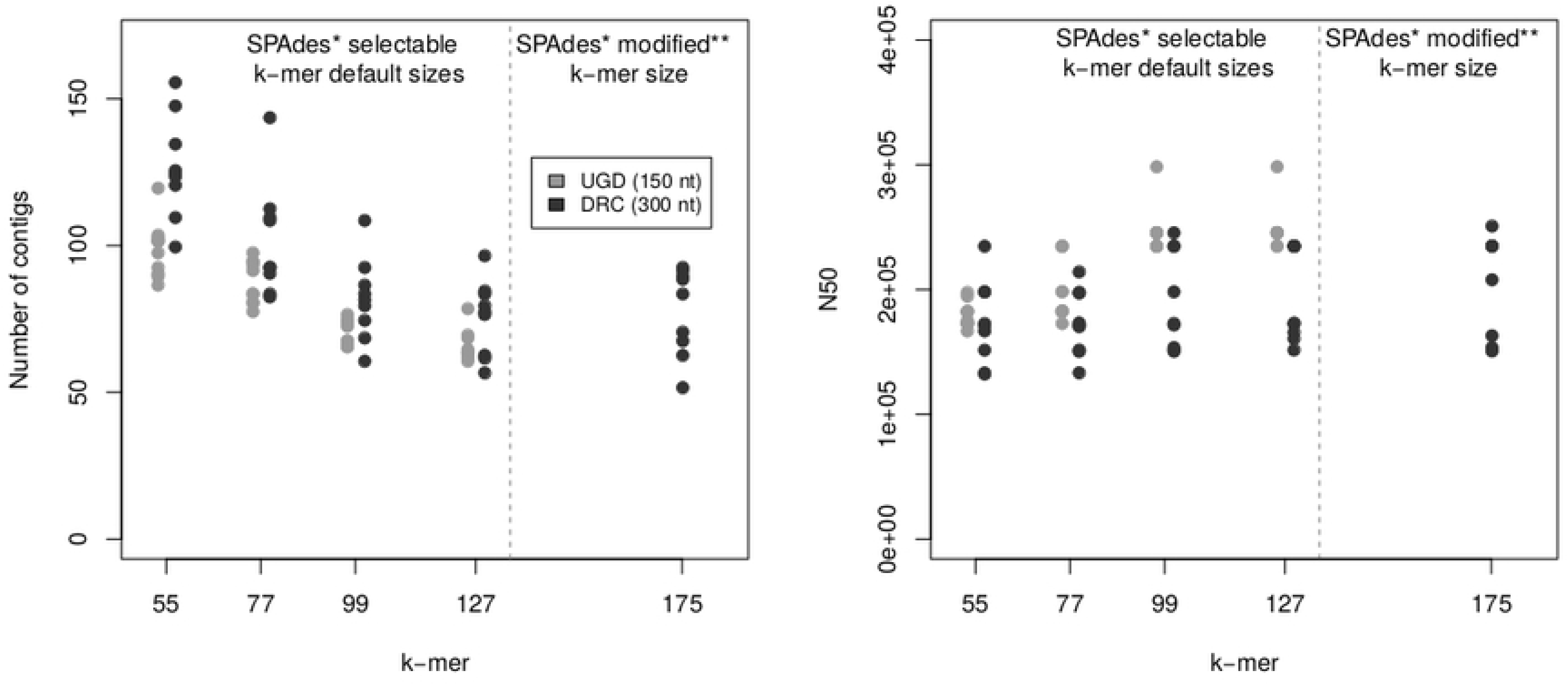
Quality metrics of genome assemblies. Number of contigs and N50 metrics, as reported by QUAST for genome assemblies from DRC and Ugandan (UGD) isolates and various selectable default and modified k-mer sizes. *: SPAdes v.3.13.0. **: After modification of the SPAdes v.3.13.0 source code.

### Comparison of analytical reagent costs

Reagents costs for MLVA typing (5 motifs) of *V. cholerae* isolates were compared using three different methods (Table 5).

**Table 5.** MLVA typing based on 5 motifs: comparative costs

|  | Cost USD |  |  |
| --- | --- | --- | --- |
|  | Sanger-based<br>(5 Motifs) | Amplicon-size-<br>based (5 Motifs) | WGS-based |
| PCR | 50 (5*10) | 50 (5*10) | - |
| Analytical cost | 35 (5*7) | 17 (5*3.4) | 150 |
| **Total** | **85** | **67** | **150** |

### Theoretical feasibility of MLVA typing using *MLVAType* on other bacterial species

As illustrated for *V. cholerae* MLVA typing, the proportion of censored estimation is directly proportional to both k-mer and VNTR length. The proportion of expected censored data was assessed *in silico* using the number of motif repeat reported in the MLVAbank for Microbes Genotyping” (http://microbesgenotyping.i2bc.paris-saclay.fr/) with a postulated 6-nt motif length and three k-mer sizes (99, 127, and 175) used during genome assembly (Fig 5).

**Fig 5.**
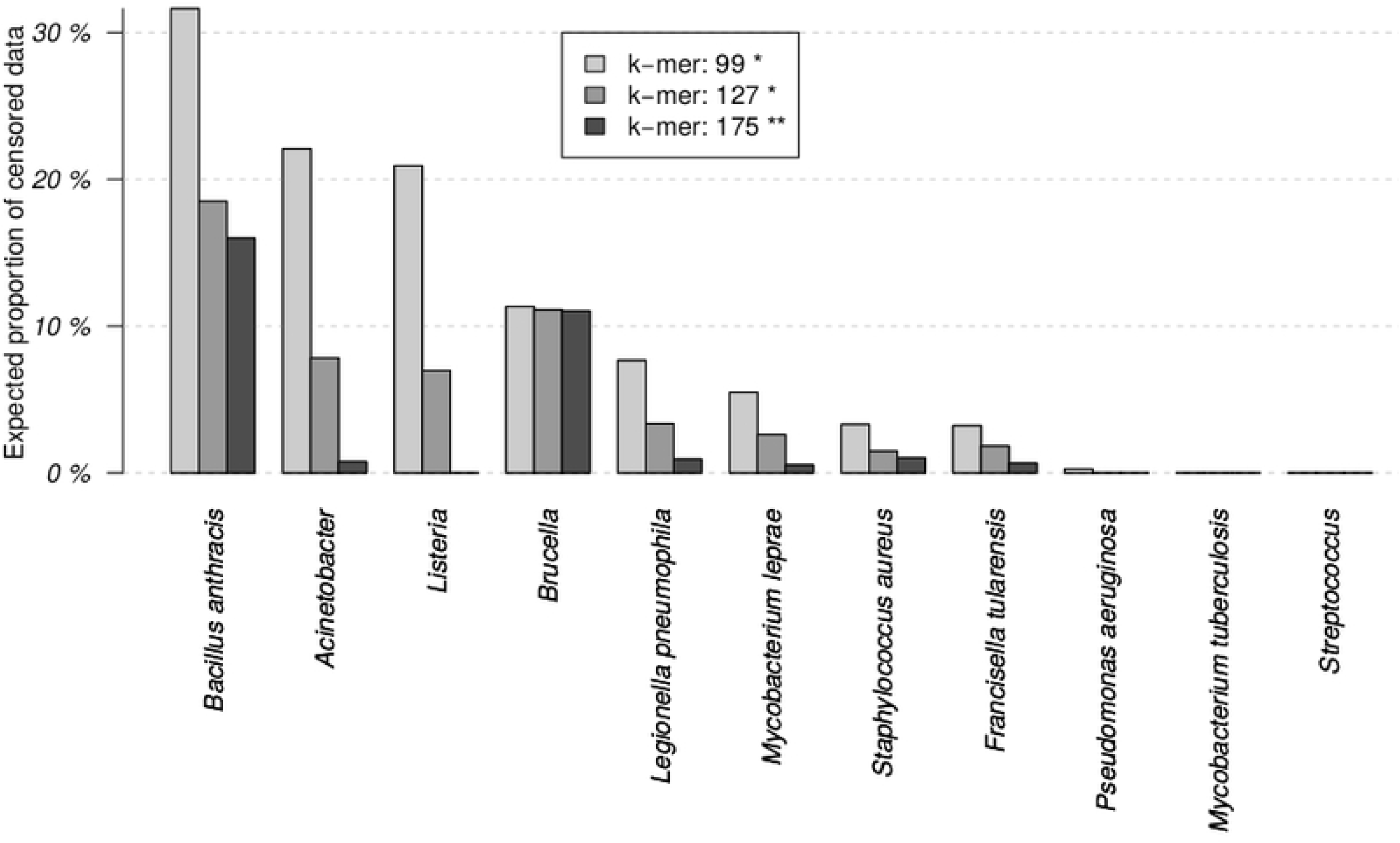
*In silico* estimation of the percentage of censored data generated by *MLVAType* application. These estimations are computed on a range of pathogenic bacteria from the MLVA Microbes Genotyping Database. *: Default sizes that are selectable in SPAdes v.3.13.0. **: After modification of the SPAdes v.3.13.0 source code.

## Discussion

The rationale behind this study is the dramatic increase in the rate and amount of sequencing and a continuous decrease in sequencing costs. The efficiency in microbial identification and subtyping allowed by WGS characterization makes it now a reference method with a clear potential to replace traditional typing methods. Anyhow, this type of analysis introduces new challenges, *i*.*e*. data storage, computing power, bioinformatics expertise and results returned in real-time, all potentially related to high costs. Undoubtedly, the WGS analysis is not yet affordable in all institutions or networks in charge of pathogen surveillance.

Accordingly, we present here a new application enabling users to extract MLVA profiles from WGS data while preserving their backward compatibility with conventional MLVA typing. To validate this application, MLVA profiles generated by *MLVAType* were compared to amplicon-size and Sanger-derived MLVA profiles on nineteen *V. cholerae* isolates among which 9 from DRC and 10 from Uganda, and the respective costs were assessed.

MLVA profiles obtained with the amplicon size-derived methods did not consistently match those obtained with the Sanger reference method and, among the former methods, Bioanalyzer results compared negatively with GeneScan data. However, it should be noted that a modification of the formula proposed by Kendall *et al*. [9], *which determines V. cholerae* VNTR repeat numbers, decreased the differences, especially with VC0437 and VCA0283, and this applied both to Bioanalyzer and GeneScan results. Bioanalyzer results could also benefit of a systematic bias correction for fragment size estimation but if building and using a conversion table, as previously proposed for *Brucella* typing [11].

In contrast and interestingly, there was no mismatch between WGS- and Sanger-derived MLVA profiles. However, WGS-derived profiles were affected by censored estimations whose proportion varied according to the k-mer size used during genome assembly with SPAdes: the larger, the k-mer size, the better the accuracy of WGS-derived MLVA profiles. Accordingly, it is recommended to use the largest possible k-mer size to assemble the reads into contigs before determining the number of tandem repeats, but considering that this maximum value is inherent to the WGS read length, as illustrated with the current application, using WGS data from *V. cholerae*. While using a k-mer size of 175 was inapplicable with read length shorter than 175, as for Ugandan isolates, it did not produce censored data with 300 nt reads from DRC isolates.

While WGS read lengths and qualities increase, one may expect to increase k-mer sizes to perform in silico MLVA typing using this *MLVAType* application whenever data comparison with MLVA database is needed. As previously reported, the length of repeat motifs should not exceed 174 nucleotides for *V. cholerae*, corresponding to 29 repetitions of a 6 nt motif [12]. Accordingly, the longer k-mer size (*i*.*e*. 175) proved to generate a correct MLVA profile with no censored data.

Current validation of *MLVAtype* algorithm with *V. cholerae* genomes paves the way for further WGS-based MLVA typing of any bacterial genome, as theoretically shown with a panel of well-recognized pathogens (*e*.*g. Streptococcus, Staphylococcus, Pseudomonas*, and *Mycobacterium* species) that are all characterized by smaller numbers of tandem repeats. With these bacteria and a postulated 6 nt motif, the proportion of censored estimations obtained with a usual k-mer size of 127 is smaller than in the current *V. cholerae* application.

## Conclusion

In conclusion, the *MLVAType* shiny application proved to extract reliably MLVA profiles from WGS data, hence solving the well-recognized issue of backward compatibility with traditional MLVA typing methods. Beside *V. cholerae*, this is also already applicable on a wide range of other bacterial pathogens and more in the future when using longer k-mer sizes. Considering the wide *in silico* exploitation of WGS data, our perspective will then be to combine the extracted information related both to VNTRs and Single Nucleotide Variants (SNVs), and to calculate a single genetic relatedness index. This should further extend our understanding of the genetic relatedness of pathogens while giving us better insight into how the VNTRs evolve over time.

## Acknowledgments

The authors are grateful to the following institutions for support provided during the course of this work: Uganda Ministry of Health, DR Congo Ministry of Health, Makerere University, John Hopkins Bromberg School of Public Health, Maryland University of Medicine, and Defense Laboratory Department (DLD), Belgium. The authors would like in a special way thank the following persons for their wonderful contribution and guidance; Professor David A. Sack, Professor Christopher G. Orach, Mr. Atek. Kagirita, Mr. Mathieu Almeida, Dr. Amanda K. Debes, M/s Shan Li, Mr. JB.Voeglein, and Dr Prudence Mitangala.

